# *Leishmania* parasite arginine deprivation response pathway influences the host macrophage lysosomal arginine sensing machinery

**DOI:** 10.1101/2021.09.01.458453

**Authors:** Evanka Madan, Madhu Puri, Rohini Muthuswami, Dan Zilberstein, Rentala Madhubala

## Abstract

Extensive interaction between the host and pathogen metabolic networks decidedly shapes the outcome of infection. Infection with *Leishmania donovani*, an intracellular protozoan parasite, leads to a competition for arginine between the host and the parasite. *L. donovani* transports arginine via a high-affinity transporter *Ld*AAP3, encoded by the two genes *LdAAP3.1* and *Ld*AAP3.2. Earlier reports show that upon arginine starvation, cultured *Leishmania* parasites promptly activate an Arginine Deprivation Response (ADR) pathway, resulting in the stoichiometric up-regulation of *Ld*AAP3.2 mRNA, protein and activity. Lysosomes, on the other hand, are known to employ a specific sensor and an arginine-activated amino acid transporter, solute carrier family 38 member 9 (SLC38A9) that monitors intra-lysosome arginine sufficiency and subsequently up-regulates cellular mTORkinase activity. The present study investigates the interaction between *Leishmania* and macrophage-lysosome arginine sensing machinery. We show that infection with *L. donovani* activates SLC38A9 arginine sensing in the human monocyte like-macrophage cell line (THP-1) when grown under physiological concentrations of arginine (0.1 mM). However, supplementing the macrophage growth medium with excess arginine (1.5 mM) followed by infection led to the down-regulation of SLC38A9. Similarly, THP-1 cells infected with *LdAAP3.2* null mutants grown in 0.1 mM arginine resulted in reduced expression of SLC38A9 and mTOR. These results indicate that inside the host macrophage, *Leishmania* overcome low arginine levels by up-regulating the transport of arginine via *Ld*AAP3 and SLC38A9 signalling. Furthermore, while *LdAAP3.2* null mutants were impaired in their ability to develop inside THP-1 macrophages, their infectivity and intracellular growth were restored in SLC38A9 silenced macrophages. This study provides the first identification of regulatory role of SLC38A9 in the expression and role of *Ld*AAP3.

**Author Summary:** *Leishmania donovani,* the causative agent of kala-azar, exhibits a digenetic life cycle. Following infection of the mammalian host, promastigotes differentiate into intracellular amastigotes within the phagolysosome of macrophages. Arginine is a central point of competition between the host and the pathogen. *L. donovani* senses lack of arginine in the surrounding micro-environment and activates a unique ADR pathway, thus upregulating the expression of the arginine transporter (*Ld*AAP3). The arginine-activated amino acid transporter SLC38A9 localizes to the lysosome surface of mammalian cells and acts as a sensor that transmits information about arginine levels in the lysosome lumen to the mechanistic target of rapamycin (mTOR) kinase. In the present study, we identified the functional interaction of host SLC38A9 and parasite *Ld*AAP3 in macrophages infected with *L. donovani.* We report that host SLC38A9 upregulation is critical for enhancing and maintaining high *Ld*AAP3 levels in intracellular *L. donovani*. Our results decode crucial information regarding the molecular mechanism involved in the arginine sensing response in *L. donovani*-infected host cells. These findings increase our understanding of the interaction of signalling intermediates during *Leishmania* infection which may lead to the discovery of novel therapeutic interventions.

## Introduction

The majority of metabolites are chemically similar or identical between the host and the pathogen; therefore, the crosstalk between their amino acid (AA) metabolic pathways is of paramount importance during infection. The host is dependent on AA metabolism to support defence against pathogens, while the pathogen is known to modulate AA metabolism to suit its own needs (1). Arginine is a versatile amino acid. It is semi-essential in mammals as it is endogenously synthesized and used for the synthesis of proteins, polyamines, urea, nitric oxide, etc. (2). Arginine is essential for *Leishmania*, as it cannot be synthesized endogenously by the parasite. It is required for the synthesis of polyamines and trypanothione (3, 4).

During infection, *Leishmania* encounters macrophage defence mechanisms designed to prevent parasite invasion and block their intracellular survival, including the release of reactive oxygen species (ROS) and synthesis of cytotoxic nitric oxide (NO) by nitric oxide synthase (iNOS) (5, 6). However, for macrophages to produce adequate amounts of NO, they must import arginine from the extracellular environment(7). *Leishmania* infection promotes the expression of arginine transporters in macrophages, including the cationic acid transporter member CAT-2 (SLC7A2) (8). Invading promastigotes activate macrophage arginase I, thereby restricting the amount of arginine available for NO production (9, 10). *Leishmania* cannot synthesize arginine *de novo* and is dependent on uptake from macrophages. The parasite imports exogenous arginine *via* a mono-specific amino acid transporter (*Ld*AAP3). Thus, reduction of the host arginine pool becomes a bottleneck for *Leishmania* in infected macrophages (9), as they are locked in competition with the host for access to the diminishing arginine supply. *Leishmania* parasites need the means to sense and respond to changes in arginine availability to be able to play the arginine hunger game successfully.

The lysosome is recognized as an essential intracellular organelle involved in the AA-sensingresponse mechanism of the mTORC1 pathway (11, 12, 13). Mammalian cells express four Rag GTPases that form heterodimers, consisting of RagA or RagB in partnership with RagC or RagD (12, 14). The amino acid-responsive super-complex that includes the pentameric Ragulator complex and the multisubunit Vacuolar ATPase (vATPase) complex promotes the accumulation of Rag heterodimers on the cytoplasmic surface of the lysosome. When Rag heterodimer is active it recruits mTOR to the lysosomal membrane, thereby initiating the activation process of mTOR (14). The lysosomal membrane represents the site at which the amino acid sensing machinery converges to stimulate mTOR (15). Solute Carrier family 38 member 9 (SLC38A9), a lysosomal arginine transporter and a sensor, is a 110 amino acid-long, soluble, cytosolic N-terminal domain responsible for its interaction with Rag-Ragulator. Two Rag GTPases and a vacuolar ATPase, along with SLC38A9, transmit information about both cytosolic and lysosomal arginine content by phosphorylating mTOR kinase (16, 17, 18, 19, 20) that subsequently activates a downstream pathway involving p-P70S6K1 (17). Thus, SLC38A9, together with a complex of additional proteins, signals mTOR on arginine sufficiency in the lysosome lumen.

We have previously reported that the *Leishmania donovani* arginine sensor responds to arginine deprivation by activating a MAPK2-mediated signalling pathway, known as the arginine deprivation response (ADR), resulting in the up-regulation of expression and activity of its amino acid transporter 3 (*Ld*AAP3) (21). Significantly, ADR is also activated during macrophage infection (21, 22, 23), indicating a role for intra-phagolysosomal arginine sensing in intracellular (amastigote) parasite survival and pathogenesis. In the present study, we have investigated the involvement of the host arginine sensing pathway by which *L. donovani* can gain access to the macrophage L-arginine pool.

Although SLC38A9 and *Ld*AAP3 have emerged as two independent lysosomal arginine transporters, a regulatory link between SLC38A9 and *Ld*AAP3 remains unknown. In the present study, we identified a critical SLC38A9 pathway that is differentially modulated after *L. donovani* infection. Subsequently, intracellular *L. donovani* derived from infected THP-1 macrophages whose SLC38A9 expression was silenced maintained in 0.1 mM arginine resulted in reduced expression of *Ld*AAP3. Similarly, we observed that infection of THP-1 cells with *LdAAP3. 2* null mutants in 0.1 mM arginine resulted in reduced expression of SLC38A9. Therefore, as there exists a positive regulatory loop between *Ld*AAP3 and SLC38A9, high SLC38A9 would lead to further increase in *Ld*AAP3 expression thereby favouring arginine transport inside the parasite. This study identifies the novel interplay of *L. donovani* and macrophage-lysosome arginine sensing machinery as a means to better understand host-parasite interaction.

## Methods

### Materials

THP-1 cell line was purchased from NCCS, Pune, India. L-arginine, Concanamycin A and Torin1(4247) were purchased from Sigma-Aldrich (St. Louis, MO, USA) and Tocris Bioscience (Bristol, UK), respectively. Fetal bovine serum (FBS), OptiMEM and Lipofectamine 3000 reagent were purchased from Invitrogen, USA. siRNAs for SLC38A9 (L-007337-02-0005) and Control (D-001810-10-05) were purchased from Dharmacon, USA. PCR primers were designed by Eurofins, USA. Antibodies were purchased from the following companies: SLC38A9 (NBP1-69235) from Novus (Littleton, CO, USA); RagA (4357S), mTORC1 (2983S) and p-P70S6K1 (Ser 389) (9206S) from Cell Signaling Technology (Beverly, MA, USA); β-actin (A1978) from Sigma (Sigma-Aldrich, St Louis, MO, USA); HRP-conjugated secondary antibodies from Cell Signaling Technology (Beverly, MA, USA).

### Leishmania cell culture

*Leishmania donovani Bob* strain (*Ld*Bob strain/MHOM/SD/62/1SCL2D) (*WT-L. donovani*) promastigotes were cultured at 26°C in M199 medium (Sigma-Aldrich, USA), supplemented with 100 units/ml penicillin (Sigma-Aldrich, USA), 100 µg/ml streptomycin (Sigma-Aldrich, USA) and 10% heat-inactivated fetal bovine serum (FBS; Biowest).

### Leishmania ADR mutant strains

*Leishmania*-adapted CRISPR/Cas9 system (24) to expedite disruption of the AAP3 genes, was used to create ADR mutants as described by Goldman-Pinkovich et al (23). These knockouts were obtained from the Prof. Zilberstein group (Technion-Israel Institute of Technology, Israel). Briefly, wild type (WT) *L. donovani* was transfected (separately) with plasmids containing three 21-nt gRNA targeting the 5’ end(173) of both *AAP3.1* and *AAP3.2* CDSs. For this, Eukaryotic Pathogen CRISPR guide RNA/DNA Design Tool was used to design gRNA sequences. These were then cloned into *Leishmania*-adapted vectorpLdCN using the single-step digestion-ligation cloning protocol previously described (23) and the constructs were later transfected into mid-log phase promastigotes. Cells were grown with G418 (50 μg/ml) for four weeks post-transfection and subsequently screened for their ability to increase *Ld*AAP3 protein abundance after arginine deprivation. Clones showing an increase in AAP3 protein abundance (*Δap3G3-17*) and another clone showing no increase in AAP3 protein abundance (*Δap3D10*) after arginine deprivation, as compared to the WT were checked for the altered protein expression (23) and selected. Once obtained, these ADR mutant cell lines were grown in G418 (100 μg/ml) supplemented culture for four-six weeks and further sub-cultured.THP-1 macrophages were then later infected with late-log phase *L. donovani* promastigotes of WT, *Δap3D10* and *Δap3G3-17* in medium containing a physiologically-relevant concentration of arginine (0.1 mM).

### THP-1 cell culture and infection

THP-1 cells, an acute monocytic leukaemia-derived human cell line (202 TM; American Type Culture Collection, Rockville, MD), were maintained in RPMI-1640 (Sigma-Aldrich, USA) medium supplemented with 10% heat-inactivated FBS (Biowest, UK), 100 units/ml penicillin and 100 μg/ml streptomycin at 37°C and 5% CO_2_. Cells (10^6^ cells/well) were treated with 50 ng/ml phorbol-12-myristate-13-acetate (PMA) (Sigma-Aldrich, USA) for 48 h to induce differentiation into macrophage-like-cells before infection. Cells were washed once with phosphate-buffered saline (PBS) and incubated in RPMI medium (Sigma-Aldrich, USA) containing 0.1 mM arginine, 10% heat-inactivated FBS, 100 units/ml penicillin and 100 μg/ml streptomycin, before infection. To carry out *in vitro* infection assays, late stationary phase promastigotes (WT and ADR mutants) were used at a ratio of 20 parasites per macrophage. *Leishmania*-infected macrophages were incubated at 37°C in a 5% CO_2_-air atmosphere for 4 h to allow the establishment of infection and proliferation of intra-macrophage parasites. The cells were then washed five times with PBS to remove non-adherent extracellular parasites. After that, the cells were incubated in RPMI medium containing 0.1 mM arginine at 37°C in a 5% CO_2_-air atmosphere for 2 h, 24 h and 48 h.

### Infectivity assay

THP-1 cells (1 × 10^6^ cells/well), treated with 50 ng/ml of PMA (Sigma-Aldrich, USA) and were seeded on glass coverslips in 6-well plates for 48 h. They were infected and simultaneously treated with inhibitors as described above, and the intracellular parasite load was visualized by Giemsa staining as described previously by Pawar et al (22). At least 10 fields were counted manually for each condition to determine the average number of parasites per macrophage.

### Small interference RNA (siRNA) transfection

Macrophage-like THP-1 cells in the exponential phase of growth were plated in a 6-well plate at a density of 1 × 10^6^cells/well and were cultured overnight. They were then transiently transfected with 60 nM of siSLC38A9 (SMARTpool: ON-TARGETplus SLC38A9 siRNA; Dharmacon, USA) (18) or a non-targeting 60 nM of siControl (ON-TARGETplus Non-targeting Pool; Dharmacon, USA), according to the manufacturer’s protocol. Lipofectamine3000 (Invitrogen, USA) in RPMI media (w/o FBS) (Invitrogen, USA) was used for transfection: 500 μl was added to each well and incubated for 24h. Then, the supernatant was aspirated, and cells were further incubated in fresh complete growth medium for 12–15 h before any treatment. Confirmation of siRNA-mediated down-regulation of the target gene was evaluated by qPCR and Western blot analysis.

### Arginine stimulation

THP-1 cells grown in RPMI were washed with PBS and maintained in 0.1mM arginine containing RPMI media. These cells were then stimulated for 2 h by the addition of RPMI containing 1.5 mM arginine. For treatment with Torin1 and Concanamycin A, THP-1 cells were infected with *L. donovani* in RPMI medium containing 0.1 mM arginine followed by treatment with 250 nM Torin1 (25) and 80 nM Concanamycin A (26) for 48 h and then harvested for qPCR and Western blot analysis. For siRNA treatment, THP-1 cells were transfected with 60 nM siRNA for 24 h, following which they were infected with *L. donovani* in RPMI medium containing 0.1 mM arginine for 48 h.

### qRT-PCR for gene expression analysis

Total RNA from infected macrophages was isolated using the TRIZOL reagent (Sigma-Aldrich, USA) and its concentration was determined by Nanodrop (Thermo Fischer, USA). cDNA was prepared from two micrograms of RNase-free DNase treated total RNA using first-strand cDNA Synthesis Kit (Thermo Fischer Scientific, USA), as per manufacturer’s instructions, using random hexamer primers. The resulting cDNA was analyzed by quantitative real-time (qRT-PCR) RT-PCR (Applied Biosystems, 7500 Fast Real-Time PCR System, CA, USA) with gene-specific primers using PowerUp SYBR Green PCR Master Mix (Thermo Fisher Scientific, USA). The details of the primers (sequences and annealing temperatures) are given in **S1Table.** Thermal profile for the real-time PCR was amplification at 50°C for 2 min followed by 40 cycles at 95°C for 15 sec, 60°C for 1 min and 72°C for 20 sec. Melting curves were generated along with the mean CT values and confirmed the generation of a specific PCR product. Amplification of RNU6AP (RNA, U6 small nuclear 1; THP-1 cells) and JW (*L. donovani*) were used as internal controls for normalization. The results were expressed as fold change of control (Uninfected samples (RNU6AP) and 2 h infected cells (JW) using the 2^-ΔΔ^*^CT^*method. All samples were run in triplicates, including a no-template (negative) control for all primers used.

### Western blotting

Western blot analysis was done as described previously by Darlyuk et al., 2009 (27). Briefly, protein was isolated from the THP-1 cells by resuspending cell lysates in RIPA buffer. Before lysis, adherent macrophages were placed on ice and washed with PBS. Macrophages were scraped in the presence of RIPA lysis buffer containing 1% NP-40, 50 mM Tris-HCl (pH 7.5), 150 mM NaCl, 1 mM EDTA (pH 8), 10 mM 1,10-phenanthroline and phosphatase and protease inhibitors (Roche).After incubation, lysates were centrifuged for 15 min to remove insoluble matter.The proteins in the lysates were quantified, 80 μg of the lysate was boiled (95°C) for 5 min in SDS sample buffer and was subjected to electrophoresis on a 10% SDS-polyacrylamide gel. Proteins were then transferred onto nitrocellulose (NC) membrane using an electrophoretic transfer cell (Bio-Rad Laboratories, USA) at RT (28). The membrane was washed with 1× TBST solution three times and blocked with 5% BSA for 2 h at RT. The blocked membrane was washed in TBST solution three times. The membrane was then incubated with primary monoclonal antibodies for SLC38A9 (1:500), RagA (1:1000), mTORC1 (1:1000), p-P70S6K1 (1:1000) and β-actin (1:5000) in PBS-Tween 20 containing 5% BSA and incubated overnight at 4^ο^C. The blots were subsequently incubated with the secondary antibody conjugated to horseradish peroxidase at 1:3000 dilution in 5% PBS-Tween 20 for 2 h at RT. Enhanced chemiluminescence reaction was used for the detection of the blot. The results were expressed as fold change and quantitated by using AlphaEaseFC image analysis software (Alpha Innotech). The data were expressed as mean ± SD of three independent experiments, and the representative image of one experiment is shown.

### Immunoprecipitation

At the specified times following infection, for immunoprecipitations, lysates were incubated with 1-2µg of primary antibody: RagA (Cell Signalling) (16 h with rotation, 4°C) and then incubated with pre-blocked Protein G beads (30ul of 50% slurry) (Sigma, USA) for 2 h at 4°C. Beads were recovered and washed two times with RIPA lysis buffer before analysis by SDS–PAGE and immunoblotting. Input was used as a positive control.

### Immunofluoroscence

THP-1 macrophages were plated on glass coverslips, and after 48 h were infected with *L. donovani*. After 48 h of infection, Confocal Microscopy was performed as described previously by Singh et al (29) using anti-SLC38A9 (NBP1-69235, Novus), anti-RagA (D885, Cell Signaling), anti-*Ld*AAP3 (gifted by Prof. Zilberstein) and anti-LAMP1 (D491S, Cell Signaling) antibodies (overnight, 4°C, 2%BSA in PBS). Images were taken with a confocal laser scanning microscope (Olympus FluoViewTM FV1000 with objective lenses PLAPON ×60 O, NA-1.42) at an excitation wavelength of 556 nm.

### Cell viability assay

MTT assay was done to assess the effect of inhibitors on THP-1 cell viability. MTT [3-(4,5-dimethyl-2-thiazolyl)-2,5-diphenyltetrazolium bromide] dye solution (Sigma-Aldrich, USA) (5 mg MTT in 1 ml PBS) was diluted 1:10 in RPMI medium (with 0.1 mM arginine). For MTT assay, 5×10^3^ THP-1 cells were seeded in each well of 96-well flat-bottom plates. Forty-eight hours after plating, uninfected and *L. donovani*-infected THP-1 cells were treated with inhibitors as described above. After 48 h, MTT assay was performed as per manufacturers’ protocol as described by Pawar et al (22). Each experiment was done in triplicates and repeated three times.

### Statistical analysis

Fold-expression (qRT-PCR and densitometric analysis) and intracellular parasite burden were represented as mean ±SD. Each experiment was repeated three times in separate sets. Statistical differences were determined using Student’s unpaired 2-tailed *t*-test. All statistics were performed using GraphPad Prism Version 5.0 (GraphPad Software, USA). p ≤ 0.05 was considered significant [* (P<0.01 to 0.05), ** (P< 0.001), *** (P< 0.0001), ns (P≥ 0.05)].

## Results

### Expression of SLC38A9 is up-regulated during *L. donovani* infection of THP-1 cells

The first set of experiments was aimed to identify the role if any, of *L. donovani* infection on SLC38A9 in infected macrophages. THP-1 cells were infected with *L. donovani* in medium containing 0.1 mM arginine (22, 30) and harvested at the end of each time point for mRNA analysis. At 2 h, 24 h and 48 h post-infection, total RNA was extracted from infected macrophages, and the resulting cDNA was subjected to real-time PCR, using *SLC38A9* specific primers as reported in the methods section. **Fig 1** shows a time-dependent increase in the expression of *SLC38A9* in THP-1 cells after infection with *L. donovani.* Maximum expression of *SLC38A9* was observed at 48 h (∼ 5-fold, p ≤ 0.005) post-*L. donovani* infection in THP-1 cells. As seen in **Fig S1A**, the infectivity of *L. donovani* in THP-1 cells at 2 h, 24 h and 48 h post-infection was between 30–40%.

**Fig 1.**
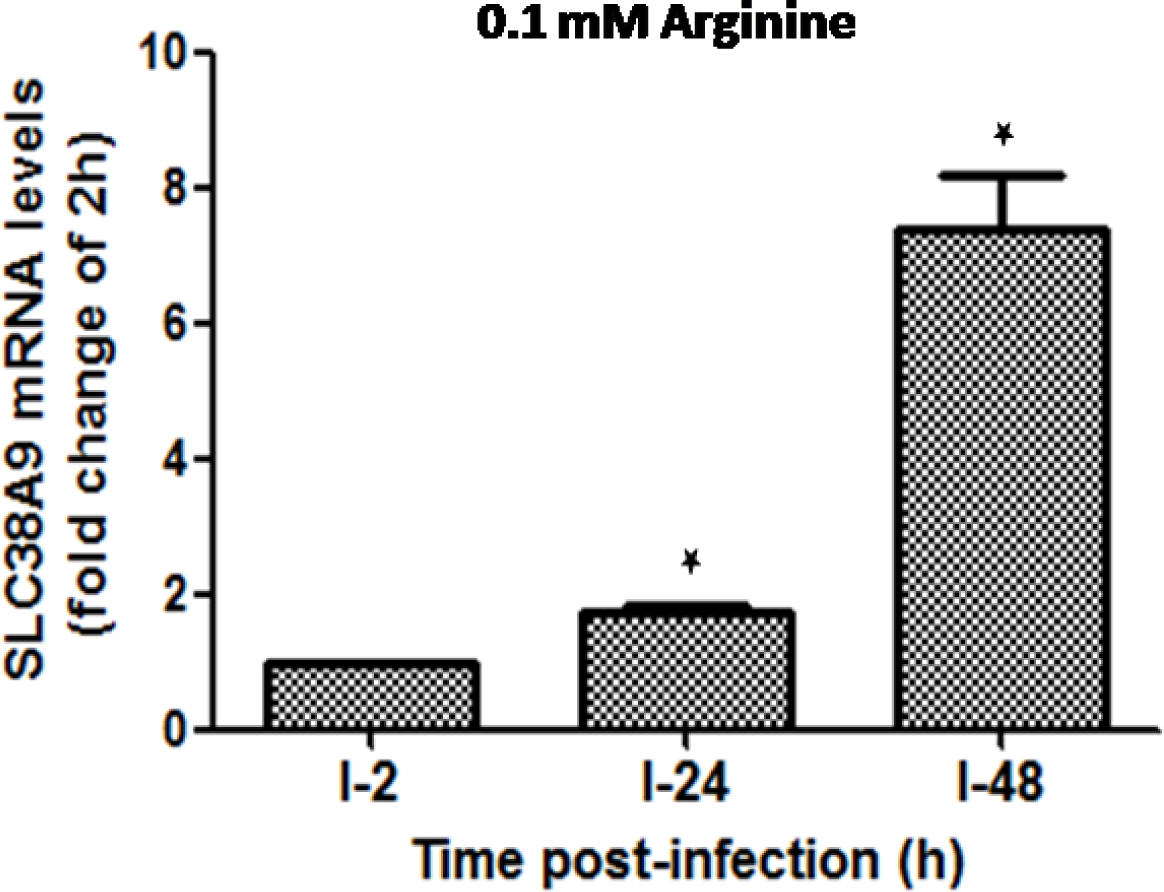
Expression pattern of SLC38A9 in *L. donovani*-infected THP-1 cells across different time points. The effect of *L. donovani* infection on the expression of SLC38A9 in THP-1 cells was analyzed. THP-1 cells maintained in RPMI medium containing 0.1 mM arginine were infected with *L. donovani* at an MOI of 20 for 2 h, 24 h and 48 h. The total RNA was extracted, and the resulting cDNA was subjected to real-time PCR analysis using primers specific for *SLC38A9.*The results are expressed as fold-change of 2 h infected cells. Data analysis was performed using the 2^-ΔΔ^*^CT^*method.Values are mean ± S.D (n = 3). The results are representative of three independent experiments performed in triplicates.

Earlier studies have shown that SLC38A9 is a lysosomal membrane protein that interacts with Ragulator and the Rag GTPases through its N-terminal 119 amino acids (‘Ragulator-binding domain’) and is required for mTOR activation (31). Several reports have indicated the role of SLC38A9 as a regulator of mTOR which further regulates protein synthesis through the phosphorylation and activation of the ribosomal S6 kinase (p70S6K1) (18, 19, 20, 32). Therefore, we investigated whether *Leishmania* infection activates host mTOR and the molecular mechanism involved in the SLC38A9 arginine sensing response in *L. donovani*-infected macrophages. For this, the expression pattern of SLC38A9, RagA, mTOR, and phospho-P70S6K1 (p-P70S6K1) (at Thr-389) was studied in infected macrophages grown in media containing either 0.1 mM or 1.5 mM arginine (22). The expression of genes was determined at the level of mRNA as well as protein in THP-1 cells infected with *Leishmania.* At 48 h post-infection, total RNA was extracted from infected macrophages, and the resulting cDNA was subjected to real-time PCR, using *SLC38A9, mTOR* and *RagA* primers. RNA derived from uninfected THP-1 cells served as a negative control. The mRNA abundance of *SLC38A9* (∼3-fold, p ≤ 0.005) and *RagA* increased (∼ 1.4 fold, p ≤ 0.005) in infected THP-1 cells cultured in 0.1 mM arginine compared to levels in uninfected (UI) THP-1 (normalized to 1) (**Fig 2A**). However, the addition of exogenous arginine (1.5 mM) to arginine deprived intracellular amastigotes resulted in the downregulation of expression of *SLC38A9* and *RagA* mRNA (**Fig 2B**).

**Fig 2.**
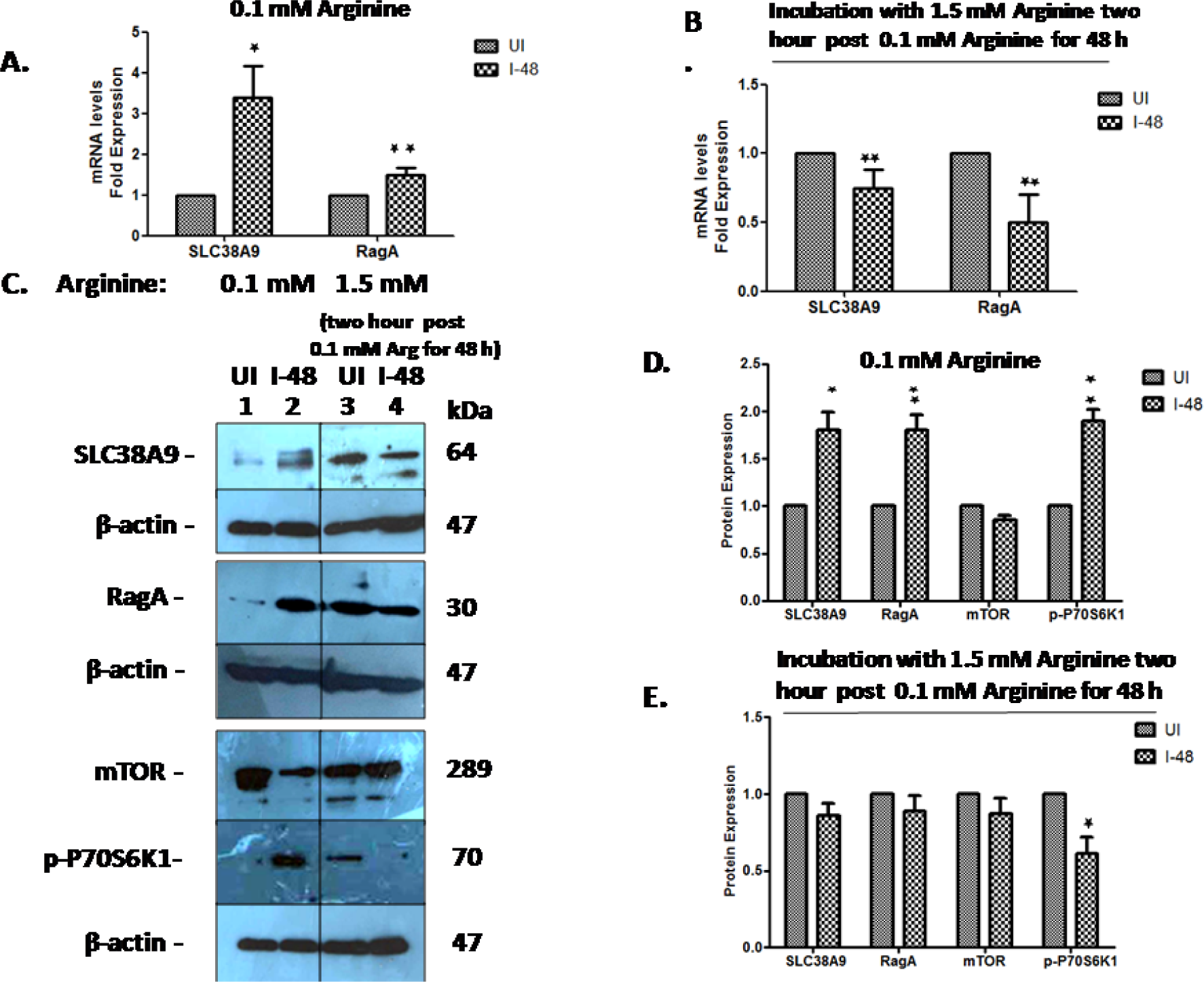
Expression pattern of SLC38A9 and mTOR in macrophages infected with *L. donovani* in different arginine concentrations. THP-1 macrophages that grew in media containing either 0.1 mMor 1.5 mM arginine were infected with *L. donovani* for 48h. (A, B)Total RNA was extracted, and the resulting cDNA was subjected to real-time PCR analysis using primers specific for *SLC38A9*and *RagA.* The results are expressed as fold-change of uninfected cells. *RNU6A* was used as a housekeeping gene. Values are mean ± S.D (n = 3). The results are representative of three independent experiments performed in triplicates. (C) Total protein was extracted and subjected to Western blot analysis using antibodies specific for SLC38A9, RagA, mTOR and p-P70S6K1. β-actin was used as a loading control. Cell lysates derived from uninfected THP-1 cultured in 0.1 mM arginine (lane 1), *L. donovani*-infected THP-1 cultured in 0.1 mM arginine (lane 2), uninfected THP-1 cultured in 1.5 mM argininetwo hours post incubation with 0.1 mM arginine for 48 h (lane 3) and *L. donovani*-infected THP-1 cultured in 1.5 mM arginine two hours post incubation with 0.1 mM arginine for 48 h (lane 4) were used. SLC38A9 levels along with its corresponding ß-actin analyzed under 1.5 mM arginine were from a different gel. The intensity of the bands was quantified by densitometry using AlphaEase FC Imager software. (D, E) The densitometric analysis shows the fold change in the expression of SLC38A9, RagA, mTOR and p-P70S6K1in THP-1 cells infected with *L. donovani*. Values are mean ± S.D. (n = 3). The results are representative of three independent experiments. (UI: Uninfected THP-1 cells, I-48: THP-1 cells infected for 48 h).

Western blot analysis using cell lysates derived from THP-1 cells was performed to confirm *Leishmania*-infection-specific regulation of the expression of SLC38A9. Cell lysates were obtained from cells cultured in 0.1 mM arginine: uninfected THP-1 (Fig 2C, lane 1), *L. donovani*-infected THP-1 (Fig 2C, lane 2). Cell lysates were also derived from cells cultured in 1.5 mM arginine: uninfected THP-1 (Fig 2C, lane 3), *L. donovani*-infected THP-1 (Fig 2C, lane 4) and were used for immunoblotting using anti-SLC38A9, anti-RagA, anti-mTOR and anti-p-P70S6K1 antibodies (**Fig 2C**). β-actin was used as a loading control. Densitometry analysis revealed a ∼2-fold increase in the levels of SLC38A9, RagA and p-P70S6K1 in *L. donovani* infected THP-1 cells as compared to those in uninfected cells when cultured in 0.1 mM arginine (**Fig 2D**). Infected THP-1 cells maintained in 1.5 mM arginine showed no significant difference in SLC38A9 and RagA levels but reduced levels of p-P70S6K1 (∼ 30%, p ≤ 0.005),indicative of reduced mTOR activity (**Fig 2E**), when compared to the levels in uninfected THP-1 cells. However, there was no significant change in the protein expression of mTOR in infected THP-1 compared to uninfected cells cultured either in 0.1 mM or 1.5 mM arginine (**Fig 2C, D and E).**These results confirm the *Leishmania* specific regulation of host SLC38A9 and mTOR activity. Additionally, the infectivity of *L. donovani* in THP-1 cells cultured in media containing different concentrations of arginine was ∼40% **(Fig S1B).**

### *L. donovani* ADR influences SLC38A9-regulated arginine homeostasis in the macrophage lysosome

We have earlier reported that the expression of *Ld*AAP3 in intracellular *L. donovani* shows a two-fold increase at 48 h post-infection when maintained in 0.1mM arginine (22).To further investigate the effect of *Ld*AAP3 on SLC38A9 arginine pathway in *Leishmania*-infected macrophages, we first examined the status of SLC38A9, mTOR expression and mTOR kinase activity (p-P70S6K1) in THP-1 cells infected with *L. donovani* CRISPR/Cas9 deletion mutant (Δ*ap3D10*) that lacks *AAP3.2* and *Ld*AAP3.2 over-expressing strain (*Δap3G3-17*) (as reported in the methods section). Δ*ap3D10* express only *AAP3.1* and do not respond to ADR, however, Δ*ap3G3-17* has increased ADR as shown by Goldman-Pinkovich*et al* 2020 (23). THP-1 cells were maintained in a medium containing 0.1 mM arginine for 48 h and were infected with either *Δap3D10* deletion mutant strain (Fig 3A, lane 2), *Ld*AAP3.2 over-expressing strain (*Δap3G3-17*) (Fig 3A, lane 3),or with WT-*L. donovani* (control; Fig 3A, lane 4).Uninfected THP-1 (Fig 3A,lane 1) cells were used as an endogenous control. Lysates from the above cells were used for immunoblotting using anti-SLC38A9, anti-mTOR and anti-p-P70S6K1antibodies. As shown in **Fig 3A and B**, there was no significant change in the expression of mTOR. However, cell lysates derived from THP-1 cells infected with the *Δap3D10* mutant showed lower levels of SLC38A9 and p-P70S6K1 (∼50-70%, p ≤ 0.005) at 48 h post-infection compared to those derived from THP-1 cells infected with WT (**Fig 3A and B**). In contrast, mutants that overexpressed AAP3 (*Δap3G3-17*) resulted in increased expression of SLC38A9 (∼1.5 fold, p ≤ 0.005) and similar expression of p-P70S6K1 in comparison to the levels in THP-1 cells infected with WT (**Fig 3A, 3B**), thereby indicating that the expression of *Ld*AAP3.2 modulates that of SLC38A9.

**Fig 3.**
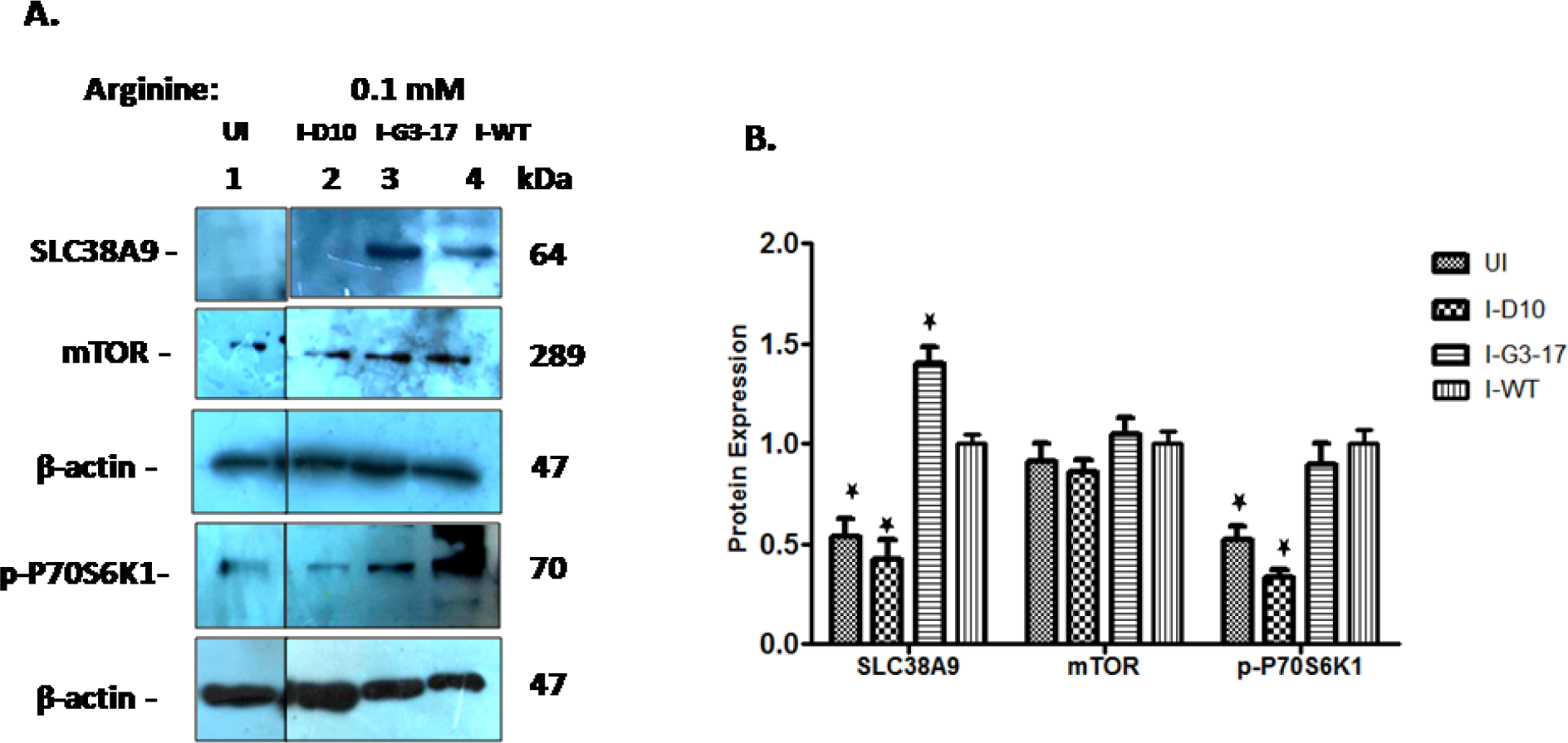
Expression pattern of SLC38A9 and mTOR in THP-1 cells infected with *L. donovani* ADR mutants. The effect of *Ld*AAP3 in the modulation of arginine sensing by SLC38A9 in *L. donovani*-infected macrophages was assessed. (A, B) THP-1 cells maintained in RPMI medium containing 0.1 mM arginine were infected with *L. donovani* deletion mutant (Δ*ap3d10*) that lacks AAP3.2, *Ld*AAP3.2 over-expressing strain (*Δap3G3-17*), or with WT-*L. donovani* at an MOI of 20 for 48 h. Protein levels of SLC38A9, mTOR and p-P70S6K1 were detected by Western blot in cell lysates using anti-SLC38A9, anti-mTOR and anti-p-P70S6K1antibodies. Cell lysates derived from Uninfected THP-1 (lane 1), THP-1 infected with either *Δap3D10* deletion mutant strain (lane 2), *Ld*AAP3 over-expressing *L. donovani* strain (*Δap3G3-17*) (lane 3), or WT-*L. donovani* strain (lane 4),in a medium containing 0.1 mM arginine for 48 h were used. ß-actin was used as a loading control. The intensity of the bands was quantified by densitometry using AlphaEase FC Imager software. The densitometric analysis shows the fold change in the expression of SLC38A9, mTOR and p-P70S6K1 in Uninfected THP-1, THP-1 cells infected with *L. donovani* deletion mutant strain, *Ld*AAP3 over-expressing strain, or WT *L. donovani*. Values are mean ± S.D. (n = 3). The results are representative of three independent experiments. (UI: Uninfected THP-1, I-D10: THP-1 cells infected with *L. donovani* deletion mutant strain, I-G3-17: THP-1 cells infected with *Ld*AAP3 over-expressing *L. donovani* strain; *Δap3G3-17*, I-WT: THP-1 cells infected with WT-*L.donovani*).

### SLC38A9 modulates ADR in intracellular parasites

Several independent reports showed that the lysosomal membrane transceptor SLC38A9 is required for arginine-mediated mTOR activation (19).We further examined the role of SLC38A9 in mTOR activation in macrophages infected with *L. donovani*. We silenced SLC38A9 in THP-1 cells by transfecting with siRNA, followed by infection with *L. donovani*. Uninfected THP-1 transfected with scrambled siRNA were used as control and normalized to 1. At 48 h post-infection, reduced mRNA abundance of *SLC38A9* of ∼90% (p≤ 0.05) was observed. These SLC38A9 transfected, infected cells expressed lower level of *mTOR* (∼30% decrease, p≤ 0.05) compared to that in control siRNA transfected cells (**Fig 4A**).Western blot analyses using cell lysates derived from THP-1 cells transfected with SLC38A9 siRNA were performed to confirm the SLC38A9-knockdown-specific regulation of expression of mTOR. As shown in **Fig 4B**, cell lysates derived from THP-1 cells infected with *L. donovani* transfected with control siRNA showed higher levels of SLC38A9, RagA, mTOR and p-P70S6K1 proteins (lane 2) (**Fig 4B, 4C**) at 48 h post-infection as compared to the uninfected control. In contrast, THP-1 cells with silenced SLC38A9 (SLCsiRNA), infected by *L. donovani* showed significant down-regulation of SLC38A9 (p≤ 0.05), RagA (p≤ 0.05), mTOR (p≤ 0.05) and p-P70S6K1 (p≤ 0.05) proteins (lane 4) (**Fig 4B, 4C**) when compared to the levels in infected cells transfected with Control siRNA (CsiRNA). This indicates thatSLC38A9 expression affects the abundance and activity of mTOR. We have previously highlighted the regulatory role of arginine in ADR modulation, as arginine starvation in intracellular amastigotes promptly activates the ADR pathway, resulting in up-regulation of the *Leishmania* arginine transporter (*Ld*AAP3) (22).To validate the role of SLC38A9 in regulating the expression of *Ld*AAP3, THP-1 cells transfected with SLC38A9 siRNA were infected with *L. donovani* in medium containing 0.1 mM or 1.5 mM arginine. At 48 h post-infection, total RNA was extracted from transfected macrophages, and the resulting cDNA was subjected to real-time PCR using *LdAAP3* specific primers. The mRNA abundance of ADR-induced *AAP3* up-regulation decreased in intracellular parasites derived from SLC38A9 siRNA transfected THP-1 cells (∼40% decrease, p≤ 0.05) (**Fig 4D**).These experiments demonstrate the regulatory role of SLC38A9 on the expression of mTOR and *Ld*AAP3. In a control experiment performed in 1.5 mM arginine, we observed a further decrease in the expression of *LdAAP3* in THP-1 cells infected with *L. donovani* after transfection with SLC38A9 siRNA (**Fig 4D**).Thus, our results indicate that ADR activation in intracellular *L. donovani* is dependent on macrophage SLC38A9 arginine sensing response.

**Fig 4.**
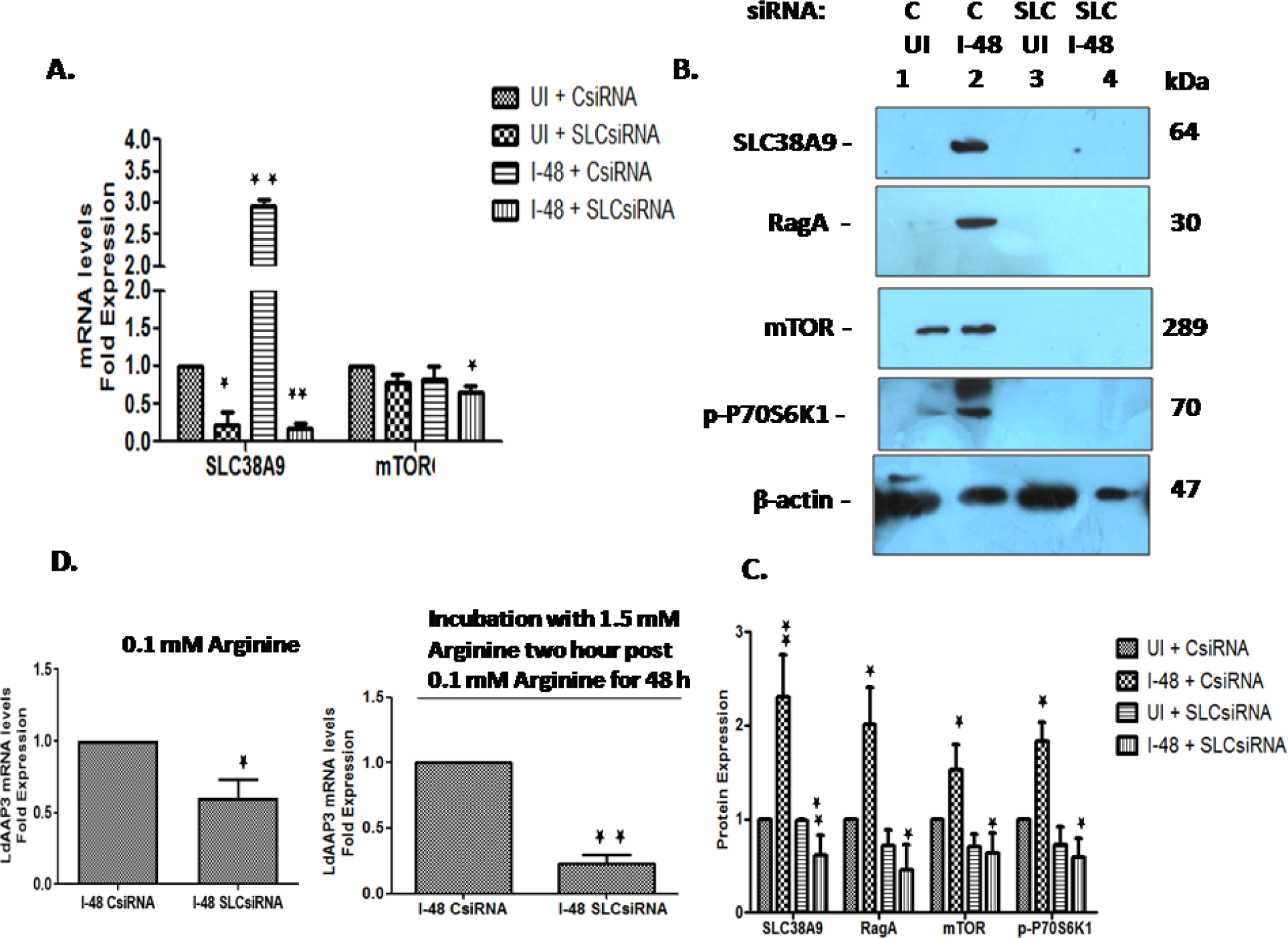
Silencing host SLC38A9 influences ADR activation in intracellular parasites. The effect of SLC38A9 in the modulation of mTOR and *Ld*AAP3 in *L. donovani* infected macrophages was checked. THP-1 cells grown in RPMI medium containing 0.1 mM arginine were transfected with 60 nM SLC38A9 siRNA for 24 h followed by infection with *L. donovani* at an MOI of 20 for 48 h. Uninfected cells transfected with Control siRNA were used as corresponding controls and set to 1. (A)Total RNA was extracted and cDNA was subjected to real-time PCR analysis using primers specific for *SLC38A9* and *mTOR.* The results are expressed as fold-change of uninfected, control siRNA transfected cells. Here, UI + SLCsiRNA, I-48 + CsiRNA and I-48 + SLCsiRNA were compared to UI + CsiRNA. *RNU6A* was used as the housekeeping gene. Values are mean ± S.D. (n = 3). The results are representative of three independent experiments performed in triplicates. (B)The total protein was extracted and subjected to Western blot analysis using antibodies specific for SLC38A9, RagA, mTOR and p-P70S6K1. Cell lysates derived from uninfected THP-1 (lane 1), *L. donovani* infected THP-1 (lane 2),uninfected THP-1 transfected with SLC38A9 siRNA (lane 3), or THP-1 transfected with SLC38A9 siRNA followed by *L. donovani* infection (lane 4), were used. Immunoblotting was performed using anti-SLC38A9, anti-RagA, anti-mTOR and anti-p-P70S6K1antibodies. Here, UI + SLCsiRNA, I-48 + CsiRNA andI-48 + SLCsiRNA have been compared to UI + CsiRNA. The intensity of the bands was quantified by densitometry using AlphaEase FC Imager software. ß-actin was used as a loading control. (C)The densitometric analysis shows the fold change in expression of SLC38A9, mTOR, p-P70S6K1, and RagA in THP-1 cells transfected with SLC38A9 siRNA followed by infection with *L. donovani*. Values are mean ± S.D. (n = 3). The results are representative of three independent experiments. (UI: Uninfected THP-1 cells, I-48: THP-1 cells infected for 48 h, C: Control, SLC: SLC38A9).(D)Bar diagrams showing fold expression of *Ld*AAP3 in SLC38A9*-*transfected infected THP-1 cells as compared to 2 h infected, untransfected control. THP-1 cells grown in RPMI medium containing either 0.1 mM or 1.5 mM arginine were transfected with 60 nM of SLCsiRNA for 24 h followed by infection with *L. donovani* for 48 h. Infected cells transfected with Control siRNA were used as controls and set to 1. The expression pattern was assessed by quantitative real-time qRT-PCR. The qRT-PCR data shown represent mean values obtained from three independent experiments. *JW* was used as the housekeeping gene. Data analysis was performed using the 2^-ΔΔ^*^CT^*method.

We further assessed whether increased *Ld*AAP3 and SLC38A9 expression are essential for infectivity and virulence in *L. donovani*. Investigations were carried out to determine the parasite load in SLC38A9 transfected THP-1cells infected with two different AAP3 mutants: *Δap3D10* and *Δap3G3-17* at an MOI of 20:1. Control siRNA transfected THP-1 cells infected with WT were used as controls. In the present study, the intracellular parasite burden was determined at 48 h post-*L. donovani* infection. WT *L. donovani* infected ∼ 50% of THP-1 macrophages, while *L. donovani* deletion mutant*(Δap3D10)* had reduced infectivity with only ∼30% macrophages (p≤ 0.05) being infected (**Fig 5A**) and *Ld*AAP3 over-expressing *L. donovani* strain (*Δap3G3-17*) showed infection comparable to that of WT parasites in control siRNA treated THP-1 cells (**Fig 5A**). Upon comparing the parasite load of the *Δap3D10* mutant in THP-1 cells transfected with control siRNA, it was observed that *Δap3D10* had 50% reduction in the number of parasites (amastigotes/macrophage) as compared to the cells infected with WT *L. donovani* (**Fig 5B**), while the parasitaemia of the *Ld*AAP3 over-expressing *Leishmania* strain (*Δap3G3-17*) was similar to that of WT (**Fig 5B**). Thus, the slow growth phenotype of the ADR mutants compared to the wild-type may be partially attributed to the reduced infectivity which is in agreement with the findings reported by Goldman-Pinkovich *et al* (23).

**Fig 5.**
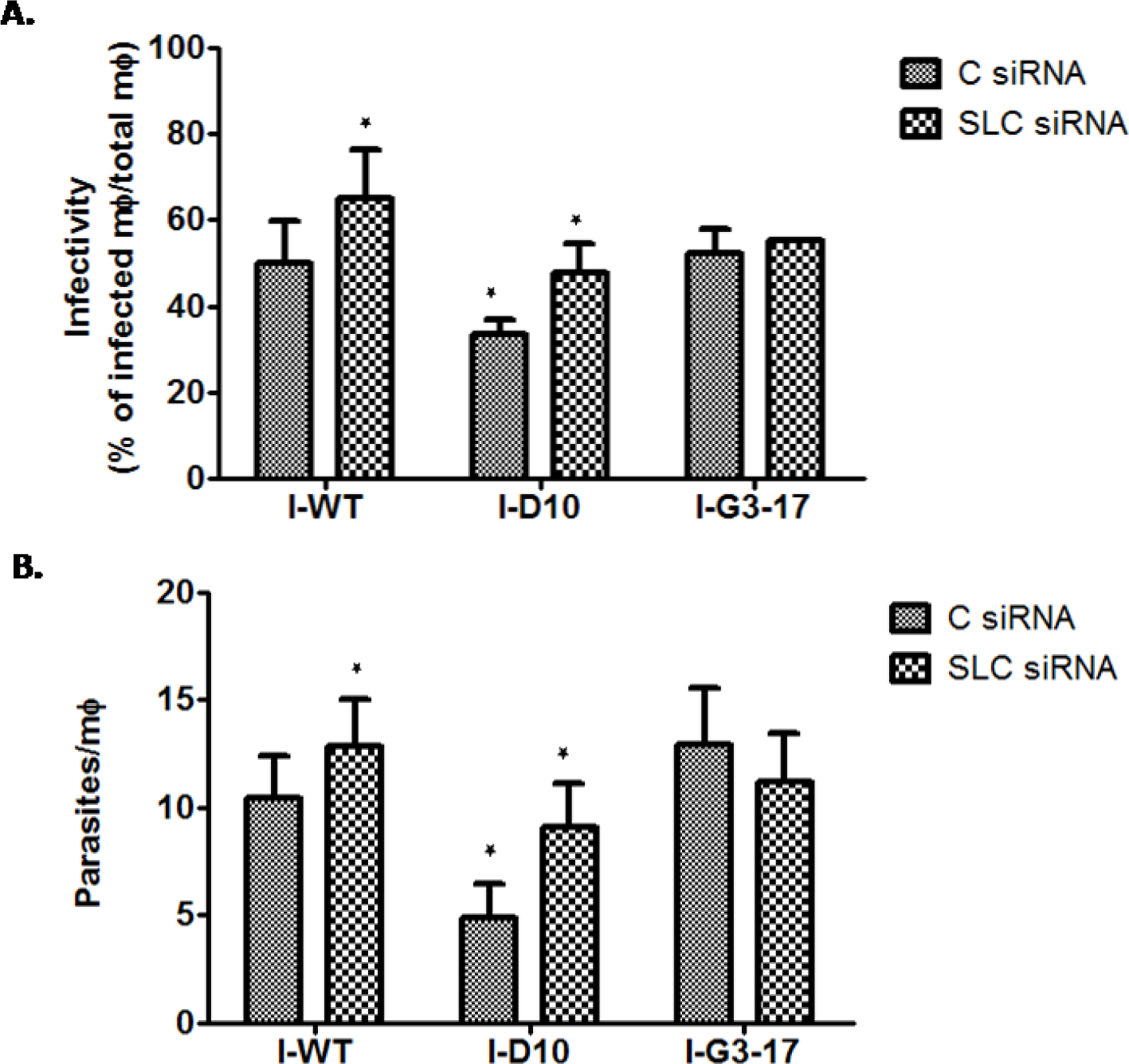
Silencing SLC38A9 in THP-1 cells increases the susceptibility of *ΔAAP3* mutants to infection. THP-1 cells grown in RPMI medium containing 0.1 mM arginine were transfected with 60 nM SLC38A9 siRNA or 60 nM control siRNA for 24 h followed by infection with *L. donovani* promastigotes of WT, *Δap3D10* and *Δap3G3-17* at an MOI of 20:1 for 48 h. (A, B) After 48 h of infection, Giemsa staining was performed to assess the parasite burden. THP-1 cells were fixed, Giemsa stained and amastigotes were counted visually. Virulence capacity was determined by calculating infectivity (A, top panel) and parasitemia (B, bottom panel).THP-1 cells transfected with control siRNA and infected with WT-*L.donovani* were used as control. The results are representative of three independent experiments. The results signify mean ± S.D with n = 3, *P < 0.05 statistical difference from the wild-type control.

Interestingly, the macrophages that lacked *SLC38A9* expression were significantly more susceptible to infection (**Fig 5A**).Infection by WT and *Δap3D10* was higher in SLC38A9 siRNA transfected macrophages as compared to that in control siRNA transfected macrophages. Consequently, it was also seen that WT and *Δap3D10* had higher parasitemia (amastigotes/macrophage) in SLC38A9 siRNA transfected macrophages as compared to control siRNA transfected macrophages (**Fig 5B**).These experiments indicate that SLC38A9 expression influences parasite ADR in a way that the mutants can resume infectivity in cells with silenced SLC38A9, grown in 0.1 mM arginine, because of retention of high levels of cellular arginine inside these cells. This points towards the role of induced SLC38A9 favouring intracellular parasite growth and infection.

### SLC38A9-vATPase complex inhibition affects ADR activation in intracellular *L. donovani*

Concanamycin A is a Vacuolar-type ATPase (V-ATPase) inhibitor (33). To assess the effect of Concanamycin A on ADR, we first examined the status of SLC38A9 and mTOR in THP-1 cells infected with *L. donovani* followed by treatment with 80 nM Concanamycin A. Uninfected and Concanamycin A-untreated cells were used as controls. At 48 h post-treatment, total RNA was extracted from Concanamycin A treated macrophages, and the resulting cDNA was subjected to real-time PCR, using *SLC38A9* and *mTOR* primers. **Fig 6A** shows a decrease in the expression of *SLC38A9* (80% reduction, p≤ 0.05) and *mTOR* (50% reduction, p≤ 0.05) in Concanamycin A treated infected THP-1 cells as compared to the levels in uninfected and untreated THP-1.

**Fig 6.**
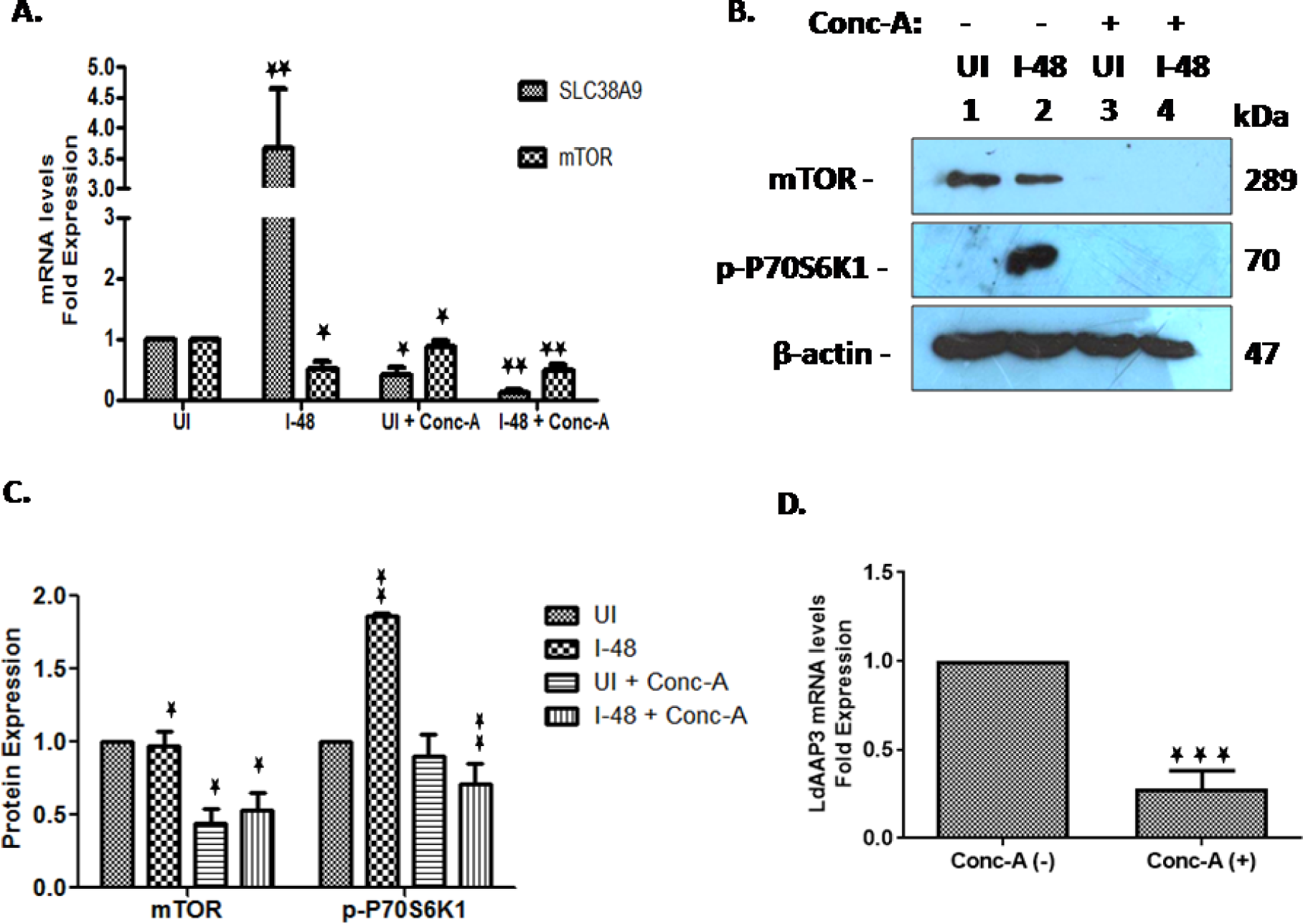
Effect of Concanamycin A on the mTOR activity in infected THP-1 cells and *Ld*AAP3 expression in intracellular *L. donovani*. The effect of SLC38A9 in the modulation of *Ld*AAP3 in *L. donovani* infected THP-1 cells was checked. THP-1 macrophages were infected with *L. donovani* promastigotes followed by treatment with 80 nM Concanamycin A for 48 h. (A) Total RNA was extracted and the resulting cDNA was subjected to qRT-PCR analysis using primers specific for *SLC38A9* and *mTOR*. Uninfected and Concanamycin A untreated cells were used as corresponding controls. The results are expressed as fold-change of uninfected and untreated cells. Here, I-48, UI + Concanamycin A and I-48 + Concanamycin A have been compared to UI. *RNU6A* was used as the housekeeping gene. Data analysis was performed using the 2^-ΔΔ^*^CT^* method. Values are mean ± S.D (n = 3). The results are representative of three independent experiments performed in triplicates. (B) Protein levels of mTOR and p-P70S6K1 were detected by Western blot in cell lysates using anti-mTOR and anti-p-P70S6K1antibodies. Cell lysates derived from uninfected THP-1 (lane 1), *L. donovani*-infected THP-1 (lane 2), uninfected and Concanamycin A treated THP-1 (lane 3), or *L. donovani*-infected and Concanamycin A treated THP-1 (lane 4), were used. Here, I-48 – Concanamycin A, UI + Concanamycin A, and I-48 + Concanamycin A have been compared to UI – Concanamycin A. (C) The densitometric analysis shows the fold change in expression of mTOR andp-P70S6K1 in infected THP-1 cells treated with Concanamycin A. The intensity of the bands was quantified by densitometry using AlphaEase FC Imager software. ß-actin was used as a loading control. Values are mean ± S.D. (n = 3). The results are representative of three independent experiments. (D) Bar diagrams showing fold expression of *Ld*AAP3 in Concanamycin A treated infected THP-1 cells as compared to 2 h infected, untreated control. The real-time PCR analysis was carried out to determine mRNA levels of *Ld*AAP3. Untreated infected cells were used as corresponding controls and set to 1. *JW* was used as the housekeeping gene. Data analysis was performed using the 2^-ΔΔ^*^CT^* method.

Western blot analysis using cell lysates derived from uninfected THP-1 (Fig 6B, lane 1), *L. donovani*-infected THP-1 (Fig 6B, lane 2), or uninfected and Concanamycin A treated THP-1 (Fig 6B, lane 3) or *L. donovani*-infected and Concanamycin A treated THP-1 (Fig 6B, lane 4) was performed. Immunoblotting was done using anti-mTOR and anti-p-P70S6K1 antibodies. As shown in **Fig 6B and C**, cell lysates derived from *L. donovani* infected THP-1 cells showed higher levels of p-P70S6K1(∼2 fold, p≤ 0.05) and similar levels of mTOR (Fig 6B, lane 2) at 48 h post-infection as compared to the uninfected control. However, Concanamycin A treatment in both uninfected and *L. donovani* infected THP-1 cells showed a significant decrease in the p-P70S6K1 (p≤ 0.05) and mTOR (p≤ 0.05) protein levels (Fig 6B, lanes 3 and 4) when compared to the levels in uninfected and untreated cells. To further investigate the role of SLC38A9 in regulating the expression of *Ld*AAP3, we examined the status of *Ld*AAP3 expression in intracellular *L. donovani* derived from infected THP-1 cells cultured in medium containing 0.1 mM arginine. The mRNA abundance of *Ld*AAP3 decreased (72% reduction, p≤ 0.05) (**Fig 6D**) upon Concanamycin A treatment in intracellular *L. donovani.* This is in concordance with results obtained with SLC38A9siRNA.

To rule out the possibility of a loss of THP-1 cell viability during infection upon Concanamycin A treatment, THP-1 cells either uninfected or infected with *L. donovani* in medium containing Concanamycin A were subjected to MTT assay to determine their viability at 48 h. As seen in **Fig S2A,** the macrophages were 75%-80% viable in media containing Concanamycin A as compared to untreated cells, thus indicating that 80 nM Concanamycin A in intracellular amastigotes was not detrimental to macrophage viability.

We further determined if Concanamycin A-induced changes in intracellular compartments were responsible for the effect on *L. donovani* intracellular growth. For this, *L. donovani*-infected THP-1 cells were treated with Concanamycin A and subjected to an infectivity assay. The percentage of infected cells and parasite load was examined at 48 h post-infection via microscopy. The infection of THP-1 cells followed by treatment with Concanamycin A showed similar parasite load as untreated cells infected with *L. donovani* **(Fig S2B).** The present data suggests that decrease in ADR activated *Ld*AAP3 levels observed upon Concanamycin A treatment is rather due to inhibition of host cell acidic compartment carrying SLC38A9-vATPase complex not due to decrease in parasite infectivity. Thus, the remodeling of the host vacuolar system may constitute a potential strategy to control *Leishmania* intracellular growth in acidic compartments.

### SLC38A9-RagA-*Ld*AAP3 form a complex at the surface of phagolysosomes during *L. donovani* infection

To rule out the possibility that the observed crosstalk between SLC38A9 and *Ld*AAP3.2 is merely an artifact produced by altered levels of lysosomal arginine concentrations, we checked for the direct interaction between *Ld*AAP3.2 and SLC38A9 using Immunoprecipitation assay. To test this, THP-1 cells were infected with *L. donovani* in medium containing 0.1 mM arginine for 48 h and harvested for immunoprecipitation followed by immunoblotting using SLC38A9, RagA and *Ld*AAP3-specific antibodies. As shown in **Fig 7**, we found that RagA efficiently pulled down both SLC38A9 and *Ld*AAP3 in THP-1 cells infected with *L. donovani*.

**Fig 7.**
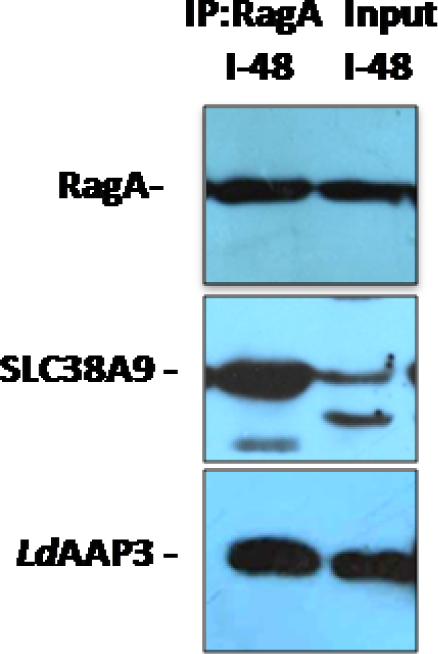
SLC38A9 interacts with RagA and *Ld*AAP3 in *L*. *donovani* infected THP-1 cells. THP-1 cells were infected with *L. donovani* in RPMI medium containing 0.1 mM arginine for 48 h. The lysates were prepared and subjected to RagA immunoprecipitation followed by immunoblotting for the indicated proteins: SLC38A9 and *Ld*AAP3. The results are representative of two independent experiments (Input: used as a positive control).

Further, the subcellular localization of SLC38A9, RagA and *Ld*AAP3 was investigated in *L. donovani*-infected THP-1 cells using confocal microscopy. Immunostaining of THP-1 cells infected with *L. donovani* for 48 h using anti-SLC38A9 and anti-*Ld*AAP3 antibodies suggested lysosomal localization of both SLC38A9 and *Ld*AAP3 as seen by their localization to TRITC-labelled lysosome associated membrane protein 1 (LAMP1)-positive vesicles **(Fig S3;panel D, E, F).** Furthermore, FITC-labelled RagA was dispersed mainly in the cytosol in uninfected macrophages, however upon infection; RagA and LAMP1 were detected together in punctuated vesicular structures throughout the phagolysosome **(Fig S3; panel E, F).** These results give direct evidence that SLC38A9 and *Ld*AAP3 directly interact at the surface of phagolysosomes and suggests that SLC38A9, RagA and *Ld*AAP3 might be part of a protein complex.

## Discussion

Induction of the arginine deprivation response in *Leishmania*-infected macrophages has been shown to play a key role in providing essential nutrients to the multiplying parasites inside parasitophorous vacuoles (PV) and/or also selectively targeting parasites to be neutralized by the innate defence response (34).Lysosomes have been recognized as a major location of the amino acid pool in mammalian cells, with sensors on the surface that report lysosome amino acid sufficiency. SLC38A9 is an arginine-activated amino acid transporter localized to the lysosome membrane of mammalian cells (18, 19). This transporter is also an arginine sensor (transceptor) (35) and together with RagGTPases recruits and activates cytosolic mTOR (18, 19, 20).Understanding the regulation of the arginine deprivation response in macrophages infected with *Leishmania* remains largely unclear. The present study firstly provides key information regarding the molecular mechanism involved in the arginine sensing response in *L. donovani*-infected host cells. Secondly, this study also reveals a novel role of SLC38A9 as a key regulator of *Ld*AAP3 in intracellular *L. donovani* and the influence of *Leishmania* ADR on host SLC38A9.

We report that the up-regulation of SLC38A9 begins at 24 h post-infection and increases five-fold at 48 h post-infection, thereby, implying that the activation of SLC38A9 in infected THP-1 cells occurs between 24–48 h post-infection. In infected phagolysosomes the initial arginine concentration is high (∼140 μM; the physiological concentration in phagolysosomes) and reduces with time (5 μM; sufficient for ADR activation in axenic *L. donovani*) (22). Subsequently, arginine concentration possibly reaches levels at 48 h post-infection, which results in the activation of SLC38A9. However, it could also be possible that induction of SLC38A9 is delayed due to the time taken to deplete intracellular parasite pools of arginine following their phagocytosis.

Arginine is a potent regulator of mTOR activity. Arginine deprivation induces eIF2α phosphorylation and decreases S6K1 phosphorylation (36, 37, 38). However, no differences have been observed in the levels of total-mTOR and phospho-mTOR (Ser 2448 and Ser 2481) upon arginine addition (37). We observed similar mTOR abundance but increased S6K1 phosphorylation in uninfected THP-1 cells maintained under 1.5 mM arginine as compared to the cells cultured under 0.1 mM arginine, as has been reported previously (36, 37). Earlier studies have shown that SLC38A9 acts on Rag GTPases and could positively regulate mTOR activity, in response to nutrient sufficiency (39, 40, 41). Furthermore, it is also known that *L. donovani* infection increases mTOR activity in mouse and human macrophages, including THP-1 cells (42, 43, 44). However, to our knowledge, no report has studied SLC38A9 mediated mTOR signalling in response to *L. donovani* infection. Thus, the present study identifies the regulatory role of *Leishmania* as the key modulator of SLC38A9-mediated mTOR pathway in macrophages maintained in different concentrations of arginine.

It is already known that following invasion, intracellular *L. donovani* parasites actively compete for the diminishing arginine pool in the phagolysosome. We have earlier reported that when arginine concentration becomes low, ADR is induced leading to the up-regulation of *Ld*AAP3 expression in intracellular *L. donovani* (22).Here, our results demonstrate a perfect temporal correlation between *Ld*AAP3 expression and modulation of SLC38A9 arginine sensing in THP-1 cells infected with *L. donovani*. We show that when *Ld*AAP3.2 is depleted in the parasites, the expression of SLC38A9 is suppressed and so is the mTOR activity; however when *Ld*AAP3.2 is overexpressed, the expression of SLC38A9 is induced. We have previously shown that wild type and *Ld*AAP3 overexpressors transport more arginine through *Ld*AAP3 (increased by ADR), while *Ld*AAP3 mutants transport normal arginine, with no further increase (23). This suggests that SLC38A9 along with *Ld*AAP3 is involved in arginine transport. Therefore, during development inside phagolysosomes, *L. donovani* may encounter a low level of arginine, and they overcome these low arginine levels by up-regulating arginine transport via *Ld*AAP3 and SLC38A9 arginine sensing. Measuring the arginine pools in lysosomes under these different circumstances in infected macrophages would be interesting follow up experiments.

A delayed response of SLC38A9 in *Leishmania*-infected macrophages suggested its possible involvement in the repression of parasite *Ld*AAP3. To investigate the biological outcome of infection-specific SLC38A9 induction, we monitored the expression level of *Ld*AAP3 in intracellular *L. donovani*. There was a marked decrease in the expression of *Ld*AAP3 in intracellular *L. donovani* derived from infected THP-1 macrophages whose SLC38A9 expression was silenced, maintained in either 0.1 mM or 1.5 mM arginine. We have previously shown that the addition of external arginine to intracellular amastigotes induces rapid degradation of *Ld*AAP3 (22). Therefore, the results of our present study point towards the cumulative effect of SLC38A9 silencing and excess arginine in reducing the levels of *Ld*AAP3,thereby indicating that the expression of AAP3 and SLC38A9 is regulated not simply by arginine levels but there exists a more direct interaction of the arginine sensing proteins/pathways.

Interestingly, it was also observed that macrophages with reduced SLC38A9 expression were significantly more susceptible to infection. Furthermore, the *Δap3D10* mutant had reduced infectivity (∼30% macrophages (p≤ 0.05) being infected) and was impaired in its ability to develop inside normal THP-1 macrophages. However, upon silencing SLC38A9 in THP-1 cells grown in 0.1 mM arginine, the ability of AAP3 mutant to develop inside these cells was found to be restored to that of the WT, because of the retention of high levels of cellular arginine inside these cells. This suggests the influence of SLC38A9 on parasite ADR. Hence, we have characterized the role of SLC38A9 during *Leishmania* infection and show that it is essential for parasite growth orinfectivity *in vitro.* Thus, inside the host macrophage, *Leishmania* must overcome the arginine “Hunger Games” by activating the SLC38A9 arginine pathway in addition to the transport of arginine via *Ld*AAP3.

The mutual influence of SLC38A9 and AAP3 prompted us to speculate that mTOR exists as a common theme between the pathways these transporters activate. To date, a role for mTOR kinase in a *Leishmania* sensing pathway has not been documented. We hypothesized that upon entry to macrophage phagolysosomes, elements in parasites arginine sensing response; MAPK2 and the ADR pathway may interact with the macrophage mTOR arginine sensing machinery that thereby induce *Ld*AAP3 up-regulation. It is known that low concentrations of Torin1 inhibit mTOR activity in THP-1 cells (45). Interestingly, in our present study it was observed that treatment with 250 nM Torin1 inhibited mTOR levels and its activity in both uninfected andinfected THP-1 cells **(Fig S4A, B and C)**.To rule out the possibility of a loss of THP-1 cell viability during infection upon Torin1 treatment, THP-1 cells either uninfected or infected with *L. donovani* in medium containing Torin1 were subjected to MTT assay to determine their viability at 48 h. As seen in **Fig S4D**, the macrophages were 80% viable in media containing Torin1 as compared to untreated cells, thus indicating that 250 nM Torin1 in intracellular amastigotes was not detrimental to macrophage viability. Thereby this indicated that under these conditions host mTOR kinases were not active. Further experiments confirming our hypothesis that the host mTOR pathway may transmit the information to parasite MAPK2 activating the PKA catalytic subunit 1 (PKA-C1) thereby leading to the up-regulation of *Ld*AAP3 expression would be planned in follow up studies.

Based on the current findings, we propose the following working model demonstrating host-pathogen interaction through SLC38A9. Upon infection in THP-1 cells, we observed increased levels of SLC38A9, RagA and a further increase in p-P70S6K1 abundance, indicative of increased mTOR activity, and consequently, increased protein synthesis (**Fig 8A**). The up-regulation of *Ld*AAP3 levels seen upon *L. donovani* infection in the previous finding (21) could be due to increased SLC38A9 levels observed in the present study, thus reflecting the regulatory role of SLC38A9 on *Ld*AAP3 via RagA/mTOR signalling. Thereby, we propose that upon infection under physiological arginine conditions, SLC38A9 transports amino acids and signals their presence at lysosomes via association with cytosolic RagA. This in turn recruits mTOR from the cytosol to the lysosomal surface. Moreover, supplementing the macrophage growth medium with arginine (1.5 mM) followed by infection with *L. donovani* results in the down-regulation of the levels of SLC38A9, RagA and p-P70S6K1, thereby possibly decreasing protein synthesis and increasing catabolism (**Fig 8B**). Our present data sheds light on the interaction between host SLC38A9 and *Ld*AAP3. Whether they interact directly or through another protein, warrants further investigation. Experiments addressing possible interactions of these proteins can be a part of follow up studies.

**Fig 8.**
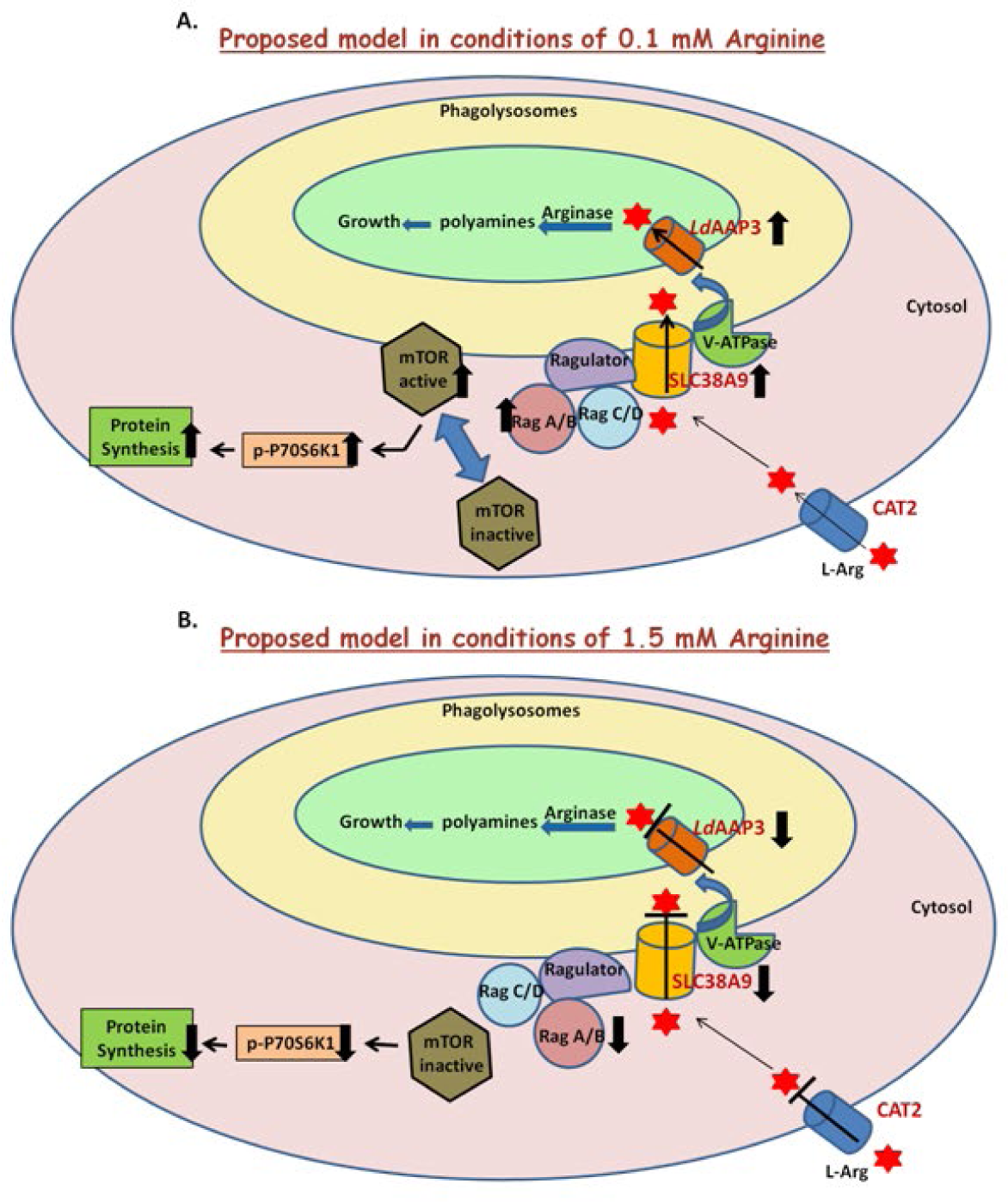
A model depicting how arginine and *Leishmania* infection signals through SLC38A9 to promote mTOR and *Ld*AAP3 activation. (A) The SLC38A9 is activated upon infection in THP-1 cellsat 0.1 mM arginine. SLC38A9-RagA complex then activates mTOR that subsequently activates p-P70S6K1. The result is up-regulation of *Ld*AAP3 expression. (B) The SLC38A9 is downregulated upon infection in THP-1 cells at 1.5 mM arginine. This then suppresses RagA and subsequently represses p-P70S6K1 and *Ld*AAP3 expression.

The present study is the first to highlight the role of SLC38A9 as a lysosomal amino acid transceptor that regulates *Ld*AAP3 expression in *L. donovani*. Our results demonstrate a link between SLC38A9 and *Ld*AAP3 and raise the possibility of SLC38A9 acting as a regulator of *Ld*AAP3, in this process. While we have studied this pathway only in macrophages, this SLC38A9-AAP3 axis may be relevant in other cell types as well.SLC38A9 could be a novel target for therapeutic intervention in leishmaniasis.

## Acknowledgements

This work was funded by a grant from the Indo-Israel Joint Research Programme (F. No. 6-7/2016(IC). Rentala Madhubala is an A. S.Paintal Distinguished Scientist Chair of ICMR and J. C. Bose National Fellow. Research fellowship to EM is from UGC D.S. Kothari (UGC, India). MP received a post-doctoral research fellowship from the Indo-Israel Joint Research Programme. We thank the Central Instrumentation Facility at the School of Life Sciences, Jawaharlal Nehru University, for providing the instrumentation facility.

