## Supplementary for "*Leishmania* parasite arginine deprivation response pathway influences the host macrophage lysosomal arginine sensing machinery"

**Running Title:** ***Leishmania*** **manipulates arginine sensing in host macrophages**

**Supplementary Data**

**Figure Legends:**

**Fig. S1: Infectivity of *L. donovani* in THP-1 cells.**

(A) THP-1 cells were infected with *L. donovani* in RPMI medium containing 0.1 mM arginine for 2 h, 24 h and 48 h. They were then stained with Giemsa and the number of infected cells was counted visually. (B)THP-1 cells were infected with *L. donovani* in RPMI medium containing 0.1 mM arginine for 48 h. They were then treated with 1.5 mM arginine for 2 h, stained with Giemsa and the number of infected cells was counted visually. The results are representative of three independent experiments.

**Fig. S2: Concanamycin A treatment does not affect the cell viability and infectivity of *L. donovani* in THP-1 cells.**

THP-1 cells grown in RPMI medium containing 0.1 mM arginine were subjected to infection with *L. donovani* at an MOI of 20, followed by treatment with 80 nM Concanamycin A for 48 h. (A) The infected THP-1 cells were incubated with MTT solution for 2 h, and stopping solution consisting of isopropanol containing 5% formic acid was added to the cells for 20 min. The absorbance was then measured at 570 nm, and the percentage of cell viability was calculated. Each experiment was done in triplicates and repeated twice. (B) The percentage of infected cells and parasite load was examined at 48 h post-infection via microscopy. The number of infected cells was counted visually after staining with Giemsa. Infected and untreated cells were used as control. The results are representative of three independent experiments.

**Fig. S3: SLC38A9, RagA and *Ld*AAP3 localize at the surface of phagolysosomes during *L. donovani* infection**

The subcellular localization of SLC38A9, *Ld*AAP3 and RagA was investigated. THP-1 cells were infected with *L. donovani* in RPMI medium containing 0.1 mM arginine for 48 h.Cells were processed and immunostained for SLC38A9, *Ld*AAP3, RagA and LAMP1. Panel A: DIC at 60X. Panel B: infected macrophages stained with DAPI. Panel C: anti-SLC38A9/ *Ld*AAP3/RagA antibody detected using FITC (green)-conjugated secondary antibody. Panel D: anti-LAMP1 antibody detected using TRITC (red)-conjugated secondary antibody. Panel E: merged micrographs and Panel F: zoomed images. (I-48: THP-1 cells infected for 48 h.

**Fig. S4: Torin1 treatment does not affect the cell viability and infectivity of *L. donovani* in THP-1 cells.**

THP-1 cells grown in RPMI medium containing 0.1 mM arginine were subjected to infection with *L. donovani* at an MOI of 20, followed by treatment with 250 nM Torin1 for 48 h. (A) The infected THP-1 cells were incubated with MTT solution for 2 h, and stopping solution consisting of isopropanol containing 5% formic acid was added to the cells for 20 min. The absorbance was then measured at 570 nm, and the percentage of cell viability was calculated. Each experiment was done in triplicates and repeated twice. (B) The percentage of infected cells and parasite load was examined at 48 h post-infection via microscopy. The number of infected cells was counted visually after staining with Giemsa. Infected and untreated cells were used as control. The results are representative of three independent experiments.

**Table. S1:** Primer sequences of the genes analyzed with the annealing temperature used for qRT-PCR

**Figures:**

**Fig S1**

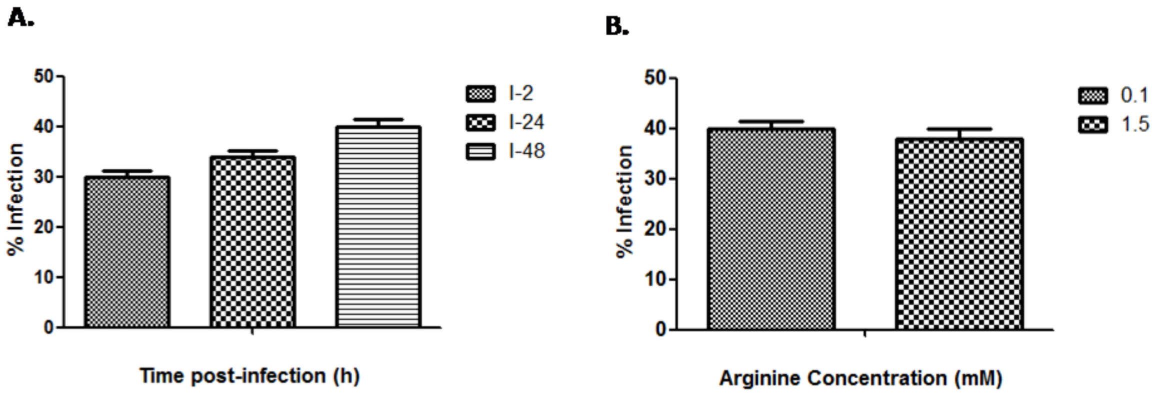

**Fig S2**

A.

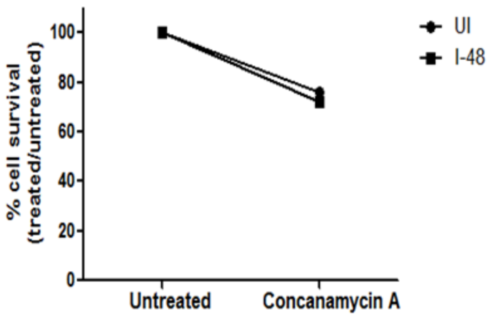
% cell survival (treated/untreated)
100
80
60
40
20
0
Untreated
Concanamycin A
UI
I-48

B.

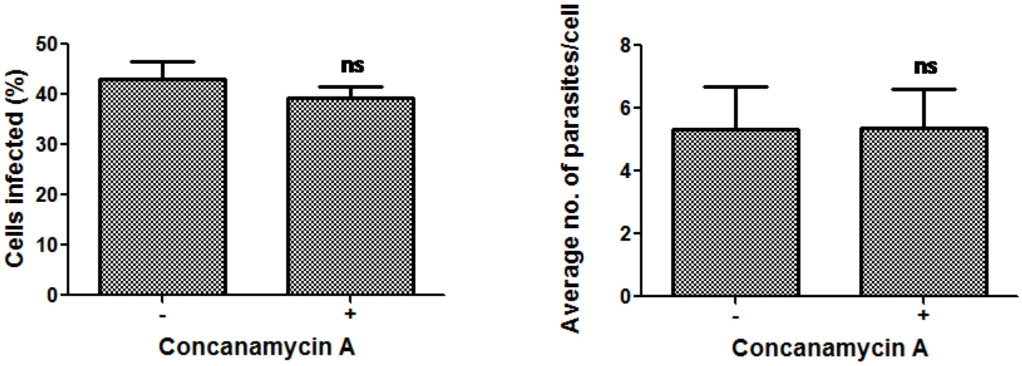
Cells infected (%)
50
40
30
20
10
0
ns
-
+
Concanamycin A
Average no. of parasites/cell
8
6
4
2
0
ns
-
+
Concanamycin A

**Fig S3**

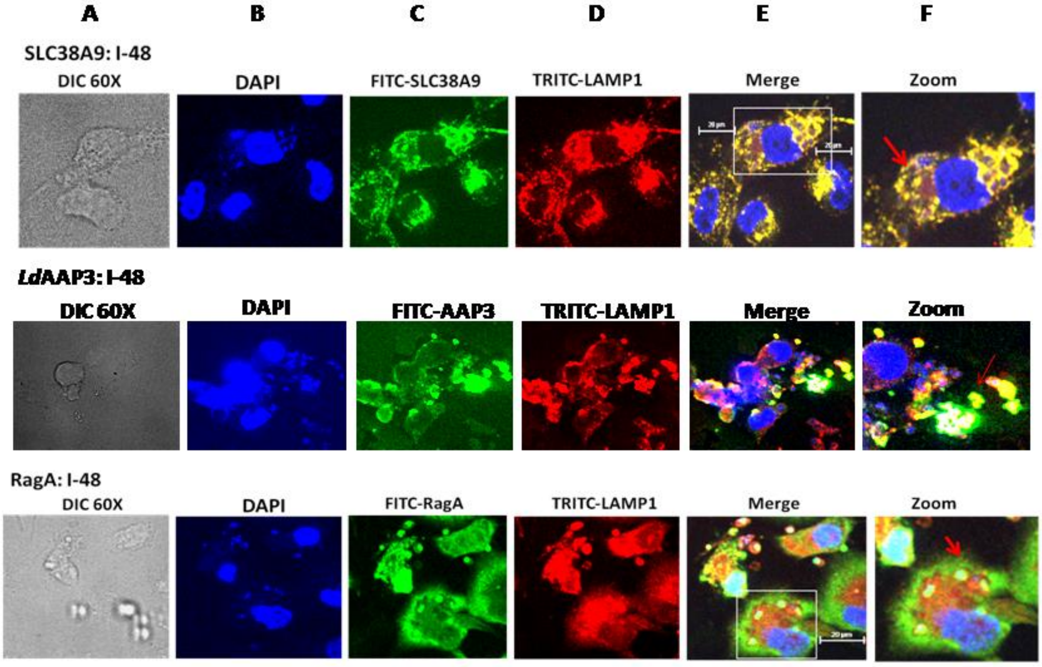

A
B
C
D
E
F
SLC38A9: I-48
DIC 60X
DAPI
FITC-SLC38A9
TRITC-LAMP1
Merge
Zoom
LdAAP3: I-48
DIC 60X
DAPI
FITC-AAP3
TRITC-LAMP1
Merge
Zoom
RagA: I-48
DIC 60X
DAPI
FITC-RagA
TRITC-LAMP1
Merge
Zoom

Fig S4

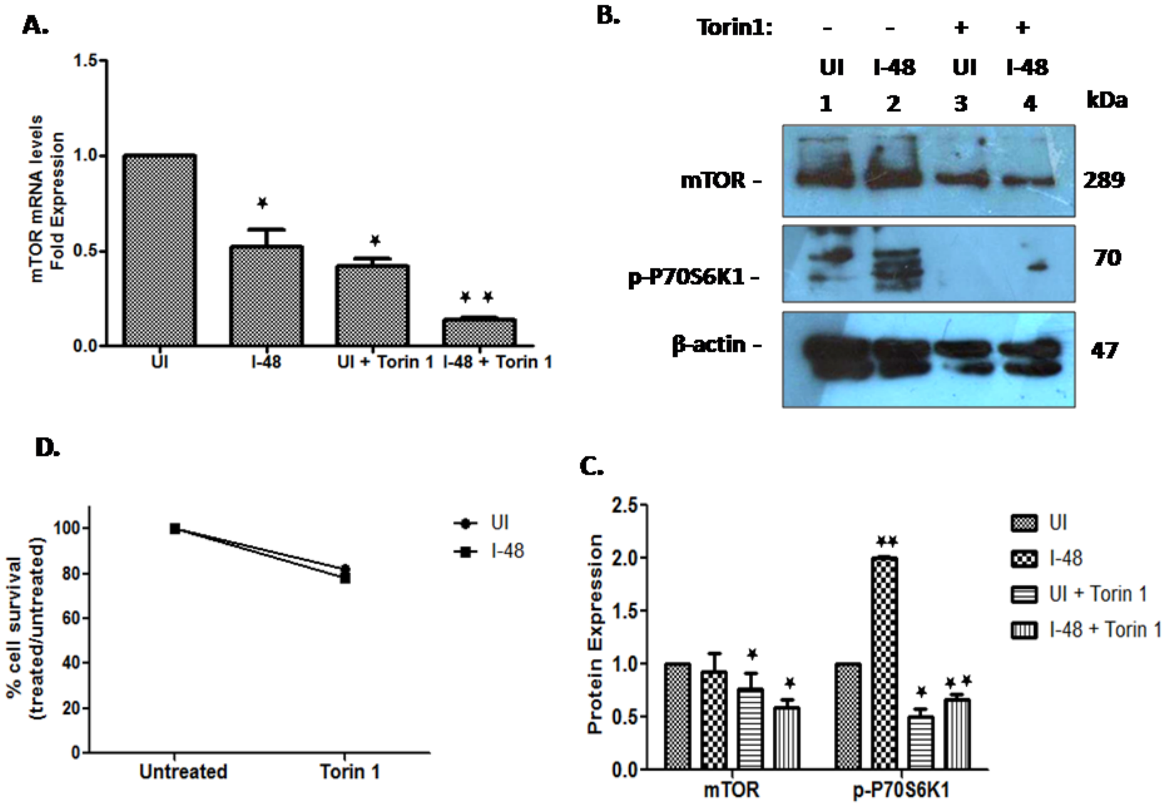
A.
mTOR mRNA levels
Fold Expression
1.5
1.0
0.5
0.0
UI
I-48
UI + Torin 1
I-48 + Torin 1
B.
Torin1:
UI
I-48
UI
I-48
1
2
3
4
kDa
mTOR
p-P70S6K1
β-actin
289
70
47
D.
% cell survival
(treated/untreated)
100
80
60
40
20
0
Untreated
Torin 1
UI
I-48
C.
Protein Expression
2.5
2.0
1.5
1.0
0.5
0.0
mTOR
p-P70S6K1
UI
I-48
UI + Torin 1
I-48 + Torin 1

### S1 Table

| Target genes | Primer sequences | Annealing temperature ($^0$C) |
|---|---|---|
| SLC38A9 | Forward 5’-TCCTTTGGGCAGTGGTCGAG–3’<br>Reverse 5’-ACTCCCGGCACTTGGACAAA -3’ | 59 |
| RagA | Forward 5’- CTCCAGAACTCTCCTGACGC-3’<br>Reverse 5’ -CTGGTAGACGATGCTGGACC -3’ | 59 |
| mTOR | Forward 5’–TCCGGCTGCTGTAGCTTATT -3’<br>Reverse 5’-CATTGTTCTGCTGGGTGAGA -3’ | 59 |
| RNU6A | Forward 5’ – CTCGCTTCGGCAGCATATAC-3’<br>Reverse 5’ –AATATGGAACGCTTCACGAATTTG -3’ | 55-60 |
| AAP3 | Forward 5’ –CGGTCGAAATGGTGCCAAAC-3’<br>Reverse 5’ –GGCTTCATCTTCCCTGCGTA-3’ | 59 |
| JW | Forward 5’ –CCTATTTTACACCAACCCAGT-3’<br>Reverse 5’ –GGGTAGGGGCGTTCTGCGAAA-3’ | 59 |
